# Negativeome in Early-Life Virome Studies: Characterization and Decontamination

**DOI:** 10.1101/2024.10.14.618243

**Authors:** Nataliia Kuzub, Alexander Kurilshikov, Alexandra Zhernakova, Sanzhima Garmaeva

**Affiliations:** Department of Genetics, University of Groningen, University Medical Center Groningen, Groningen, the Netherlands

## Abstract

Environmental contamination complicates the study of low-biomass microbial communities like the human gut virome. In 1,321 early-life gut viromes and 55 negative controls (NCs) from four datasets, we identified viral contaminants and their prevalence across studies. Samples and NCs were indistinguishable in genomic and ecological features, with 71.5% of samples sharing at least one identical strain with NCs. This work demonstrates the efficacy of strain-aware decontamination for preserving biological signals.

## Main text

Viruses are the most abundant biological entities on Earth, and they constitute a significant component of the human gut microbiome. The number of viruses in the human gut has been estimated to exceed 10^12^, roughly equal to the number of bacteria^1^. Despite this quantity, the total weight of the human gut virome’s genetic material can be just ∼50 microgram^1^. This makes virome extraction and annotation challenging, particularly in low-diversity samples like those from early life^2–4^. A major issue here is distinguishing genuine sample signals from environmental contamination, a topic of recent debate in relation to the microbiome of the placenta^5^ and blood^6^. To address this issue, proper negative controls (NCs) are essential at every step of sample processing. Best practices include sequencing NCs alongside the samples and removing sequences identified in NCs during data analysis^2–4,7,8^. However, the impact of contamination on virome samples, its sharedness across different studies, and appropriate decontamination strategies remain under-studied.

Here, we aimed to characterize the viral composition in NCs, assess the impact of environmental contamination on the samples (particularly low-biomass samples), and elucidate the level of genomic resolution necessary for virome decontamination. To do so, we employed publicly available data from four early-life virome studies^2–4,9^ that had used viral-like particle (VLP) enrichment protocols and deposited raw sequencing data for both biological samples and NCs in public archives. Together, these studies include 1,321 biological samples (1,175 infant samples from 0 to 5 years and 146 maternal samples) and 55 NCs (Supplementary Data 1, Supplementary Figure 1), in which we identified 971,583 putative virus sequences clustered to 193,970 viral operational taxonomic units (vOTUs) (Supplementary Figure 2a,b).

We first aimed to determine if specific features (number of reads and reconstructed contigs, virus sequences, viral genome completeness, and viral diversity and richness) could distinguish NCs from biological samples. However, we found no significant differences between NCs and samples for any of these features (p > 0.1, Supplementary Data 2–7, Figure 1a,b, Supplementary Figure 3). While we did observe significant differences in richness and diversity between NCs and samples in two studies when tested separately (FDR < 1e-02, Figure 1b, Supplementary Data 6 & 7, Supplementary Figure 3), the direction of association was inconsistent. The average number of vOTUs detected varied substantially across studies (intraclass correlation coefficient (ICC) = 0.2, Supplementary Data 8). Biological samples had a median of 230 vOTUs (interquartile range (IQR): 64–594), whereas NCs had a lower median of 5 vOTUs (IQR: 0–33), although this difference was not statistically significant. Overall, NCs did not differ from biological samples in any major genomic and ecological features.

**Figure 1:**
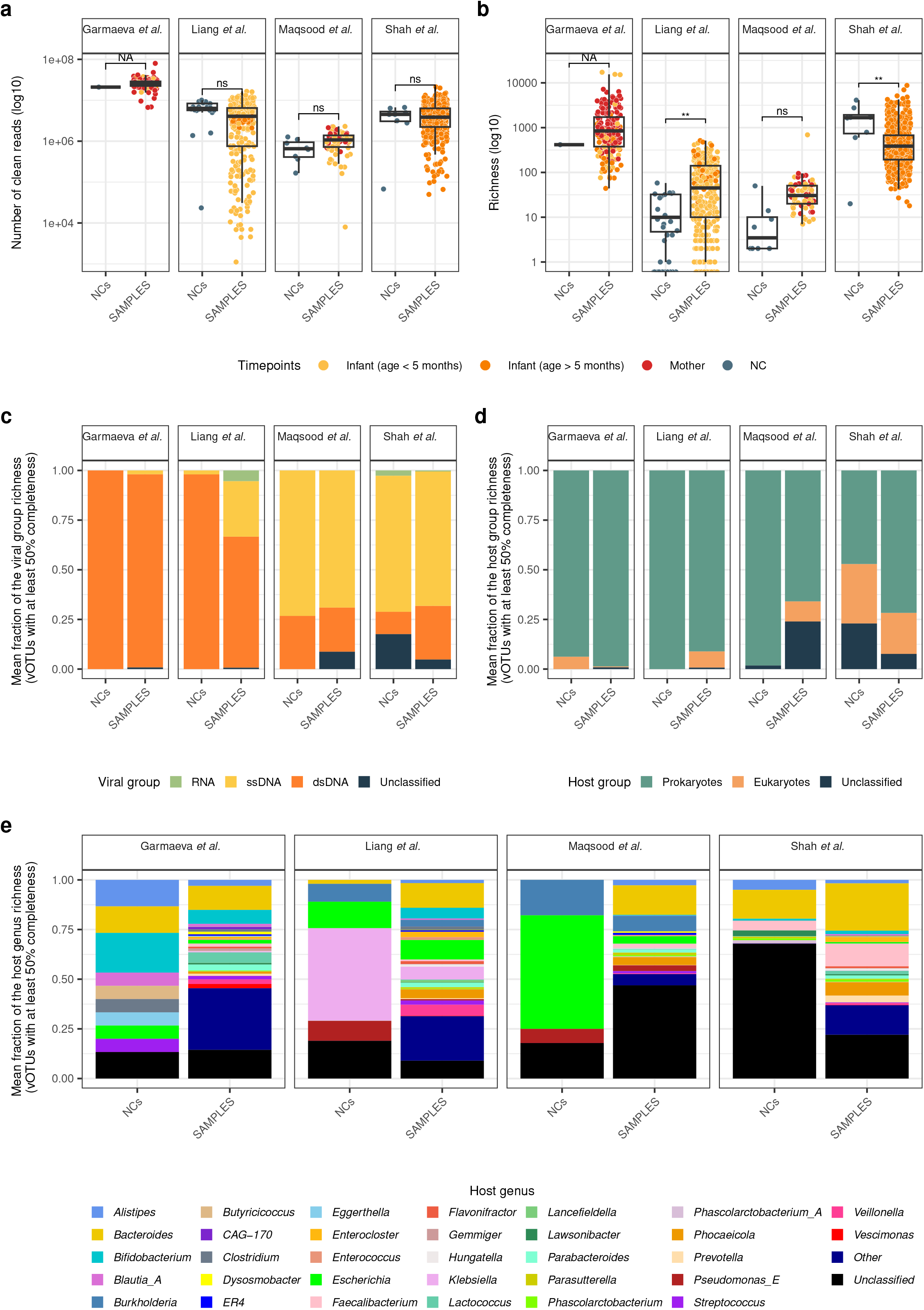
Key genomic and ecological features in negative controls (NCs) and samples. **a**. Number of clean reads in NCs vs samples. **b**. Viral operational taxonomic unit (vOTU) richness in NCs vs samples. In **a** and **b**, every dot is a sample, and color indicates age: infant samples (age < 5 months) in yellow, infant samples (age > 5 months) in orange, maternal samples in red, and NCs in dark blue. Boxplots visualize the median, hinges (25th and 75th percentiles), and whiskers extending up to 1.5 times the interquartile range from the hinges. **c**. Mean vOTU proportion by nucleic acid type assigned via deduced virus taxonomy. **d**. Mean vOTU proportion by the assigned host domain (eukaryotic vs prokaryotic). **e**. Mean proportion of the vOTUs grouped by the predicted bacterial host genus in NCs vs samples.

We next investigated differences in virome composition between NCs and biological samples, categorizing vOTUs with ≥ 50% genome completeness by nucleic acid type (dsDNA, ssDNA, RNA) based on their assigned taxonomy. Here, we observed a significantly lower proportion of ssDNA viruses in NCs compared to samples across all studies (beta = -14.9, FDR = 1e-04, Supplementary Data 9), driven mainly by a significant difference in one of the studies (Liang et al., FDR = 2e-02). When further tested separately, samples from Shah et al. had fewer RNA viruses than NCs (beta = -1.6, FDR = 2.3e-05, Figure 1c, Supplementary Data 9).

At the level of the predicted host domain, prokaryotic viruses dominated all datasets (p = 2.4e-129, Figure 1d, Supplementary Data 10), consistent with previous findings^10,11^, with no difference between NCs and samples (FDR > 0.1, Supplementary Data 11, Figure 1d). Study-specific testing revealed a lower proportion of prokaryotic viruses in NCs compared to samples in one study (FDR = 2e-05, Shah et al.). At the genus-level of prokaryotic viruses’ hosts, NCs primarily contained viruses with hosts typical of the human gut microbiota, such as *Alistipes, Bacteroides, Bifidobacterium*, and *Escherichia* (Figure 1e). These observations demonstrate that the viruses identified in NCs resemble the viruses of the human gut, at least at the highest taxonomy and predicted host levels.

We further investigated whether NCs from different studies share common viruses. Among the 5,984 vOTUs identified in NCs from all four studies, none were common across all studies. However, two vOTUs were found in NCs from three studies and 44 vOTUs were found in two studies, while 5,938 were study-specific. At strain-level (see Methods), only three viruses were shared between NCs and samples from two studies. Of note, two Caudoviricetes phages found shared between two studies from the U.S.^3,4^ are predicted to infect *Burkholderia cepacia* complex species (Figure 2a), which are common environmental contaminants often shared across U.S. hospitals^8,12,13^. Another strain-level virus shared across two studies, phiX174, is commonly used as a positive sequencing control (Figure 2b).

**Figure 2:**
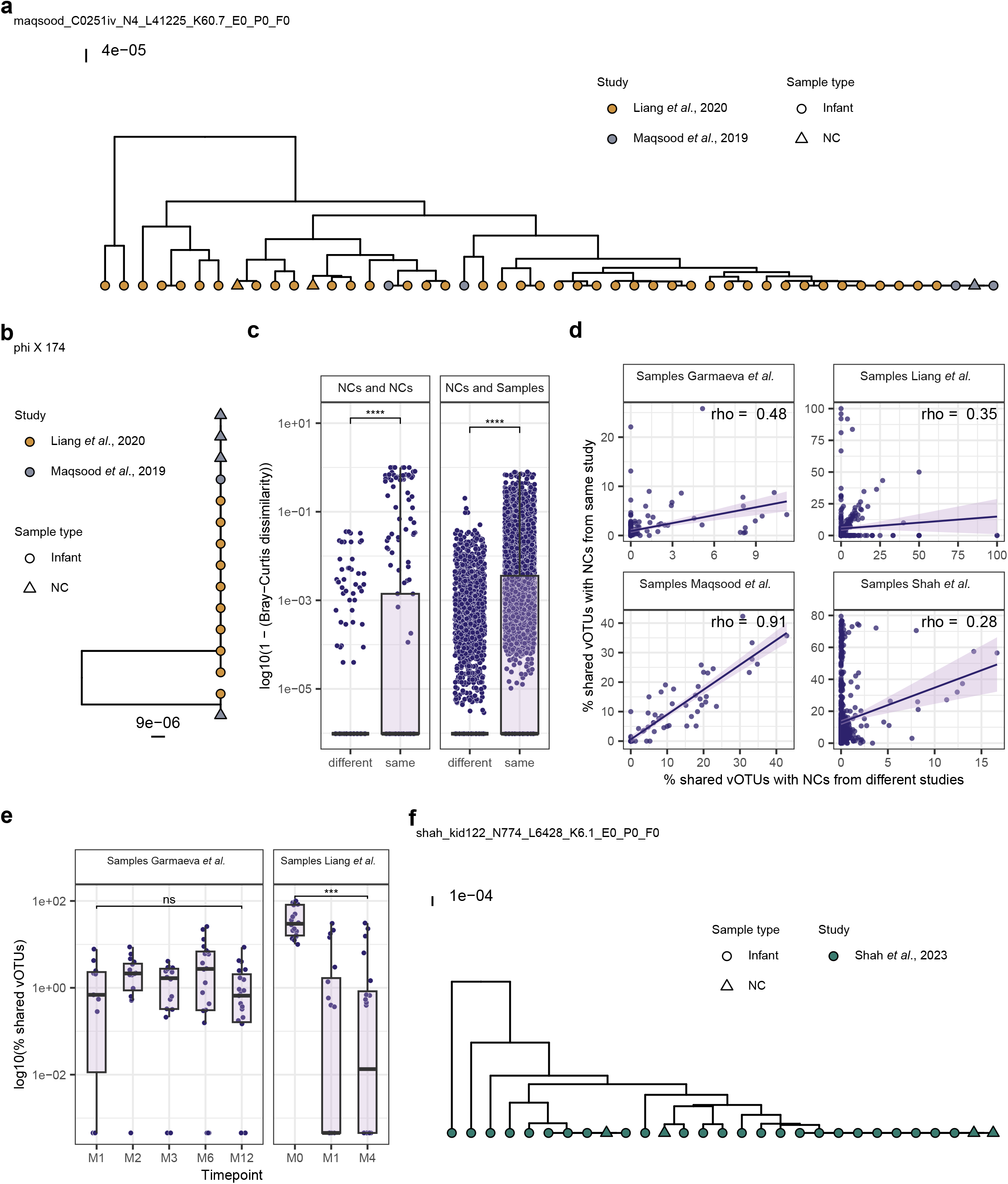
Estimation of contamination in samples and influencing features. **a**, **b**. Reconstructed ultrametric trees for the Burkholderia phage and phiX174, respectively. Every point is a sample. Point shape indicates sample type: infant (circle) and NC (triangle), and point color represents the study. **c**. Sample similarity (1 – Bray-Curtis dissimilarity index). Left vertical panel compares NCs across the different and same studies. Right panel compares NCs with samples from the different and same studies. **d**. Correlation of percentage of vOTUs shared between samples and NCs across different and same studies. The percentage of vOTUs shared with NCs is calculated as the number of vOTUs shared with NCs divided by the total sample richness. **e**. Percentage of vOTUs shared between samples and NCs per infant timepoint, separated by study. Boxplots visualize the median, hinges (25th and 75th percentiles), and whiskers extending up to 1.5 times the interquartile range from the hinges. **f**. Reconstructed ultrametric tree for a Bacteroides phage.

Overall, NC vOTU compositions were more similar within the same study than across different studies (FDR < 3e-04, Figure 2c, Supplementary Data 12). Of all the vOTUs detected in NCs per study, 7.7–23.9% were shared within the same study at strain-level. NCs were also more similar to samples within their study than to samples from other studies (FDR < 4e-04, Figure 2c, Supplementary Data 12). In general, most of the NC viromes were study-specific, but a few viral strains were shared across studies, reflecting common environmental and technical contamination.

We then quantified sample contamination by analyzing the proportion and abundance of vOTUs shared with NCs within each study. Only 20.3% (N=256) of all biological samples did not share any vOTUs with any NCs, while some shared up to 100% composition. 75.3% of the samples overlap with NCs at vOTU-level, with a median of 4.9% (IQR: 1.9–12.9) of vOTUs per sample shared with NCs from the same study, with a median abundance of 1.7% (IQR: 0.3–8.7). Moreover, 41.1% of samples shared a median of 0.7% (IQR: 0.2–4.8) of vOTUs per sample with NCs from other studies, though this vOTU-sharing was significantly lower than to own NCs in two out of four studies (Supplementary Data 13, Supplementary Figure 4a). The correlation between the percentage of vOTUs shared with the study’s NCs and those from other studies was moderate to strong (ρ > 0.2, p < 1.3e-10, Figure 2d, Supplementary Data 14). This suggests that the sample contamination level could be estimated using the proportion of sample vOTU-sharing with NCs from independent studies. To facilitate the use of available NCs in quality control of future studies, we have made the database of vOTU sequences identified in NCs publicly available^14^.

Given the lower gut viral diversity reported for infants compared to adults^2^, we investigated whether contamination levels differed between maternal and infant samples and across infant ages. In the two studies where both infant and maternal samples were available, we found that the proportion of vOTUs shared with NCs from the same study was significantly higher in infants than in mothers (beta = 2.5, FDR = 5.1e-06 in Garmaeva et al.; beta_2_ = 11.6, FDR = 2.3e-07 in Maqsood et al.; Supplementary Data 15; Supplementary Figure 4b). The estimated contamination decreased with infant age but was significant in only one of the two longitudinal studies (beta = -0.1, FDR = 0.3 in Garmaeva et al.; beta_2_ = -7.7, FDR = 2e-04 in Liang et al.; Figure 2e; Supplementary Data 16). These observations suggest that lab-induced contamination is more likely to be sequenced and detected in early-life gut virome samples due to their low diversity and low overall viral load.

Since NCs harbored viruses similar to those in the human gut at predicted host-level, we assessed whether the overlap identified at vOTU-level extends to strain-level, indicating sample contamination, or simply reflects broader taxonomic similarities between environmental and gut viromes. We therefore estimated the proportion of samples sharing identical strains with their study’s NCs. To do so, we reconstructed virus strains for 5,635 vOTUs present in NCs and biological samples (see Methods). We found that only 32.6% of these vOTUs (N=1,838) were shared between NCs and at least one biological sample at strain-level, suggesting that the strain detected in the sample might originate from environmental contamination (see Methods, Figure 2f). Across the entire dataset, 71.5% (N=806) of samples had at least one strain identical to one detected in NCs. Since more than half of vOTUs shared between samples and NCs differed at strain-level, likely representing true biological signals, we conclude that decontamination of NC-detected viruses from the dataset should be performed at strain-level.

After performing strain-level decontamination, the vOTU richness in samples sharing vOTUs with NCs dropped by 1.5% (IQR: 0.5–5.0) on average. The number of vOTUs shared with own NCs decreased by 33.3% (IQR: 14.9–50.0) and comprised 2.9% (IQR: 1.0–7.8) of all vOTU detected per sample. Next, we compared the decontamination results at strain-versus species-level. For the latter, all vOTUs shared with NCs were excluded, leading to a 4.9% (1.9–12.9) drop in vOTU richness per contaminated sample, which was significantly higher compared to the strain-level decontamination. These results further demonstrate that strain-level decontamination preserves key ecological features of the samples, such as richness.

In summary, our results show high similarity between NCs and samples in key ecological and genomic features. Although biological samples demonstrated compositional similarity to NCs, most of the viruses shared between NCs and samples represented different strains. We therefore recommend performing decontamination at strain-level rather than vOTU-level as this provides higher precision. In this study, we also provide a pipeline for strain-level decontamination.

We also observed study-specific NC composition that likely represents variation between experimental environments. However, we also observed that sample vOTU-sharedness with its NCs is correlated to sharedness with NCs in other studies, a measure that may help to estimate the level of sample contamination in study designs that lack NCs. Lastly, contamination in virome data is unavoidable, so NCs from multiple sources at different processing stages must be sequenced and deposited with samples for future data reuse. This is especially important for early-life virome studies, as we showed that infant feces sampled closer to birth are more prone to contamination and share more vOTUs with NCs than feces sampled later in life.

## Methods

### Publicly accessible datasets of VLP-enriched early-life human gut samples

Detailed descriptions and metadata of the studies mentioned below can be found in the original articles^2–4,9^. Here, we briefly summarize the set-up of each study.

*Garmaeva* et al.^2^ was based on samples collected longitudinally from 32 mother–infant pairs of the Dutch LLNEXT cohort^15^. In total, 86 VLP samples were recovered from 32 infants (including two twin pairs) during the first year of life. 119 maternal samples from 30 mothers were collected longitudinally from 28 weeks of pregnancy to 3 months postpartum. Four negative buffer controls were isolated with the samples: three failed sequencing and the other is included in the current study.

*Liang* et al.^3^ was completed in Pennsylvania Hospital, Philadelphia, USA and includes VLP data for 185 samples collected longitudinally from 144 infants from 0–4 months and 21 samples collected from 21 children from 2–5 years. For all infant samples, both DNA and RNA VLP sequencing was done. For samples of children from 2–5 years, only DNA VLP sequencing was done. We included both the DNA and RNA sequencing samples in our study and analyzed them as separate samples. The *Liang* et al. study included 19 NCs of different origins, including empty diaper samples, empty stool container samples, and reagent-only samples. For each NC, both DNA and RNA sequencing were performed, further doubling the number of NC samples to 38. While all these NCs were initially included in our study, we did not discover any putative viral sequences in 12 out of 38 NC and did not detect any vOTU in 18 out of 38 NC samples.

*Maqsood* et al.^4^ is a study conducted in St. Louis, Missouri, USA that analyzed the birth stool of twins alongside samples collected from their mothers. Samples from 51 infants and 27 mothers were available for this study. Two different types of NCs were included: buffer NCs (n=4) isolated to describe general contamination and NCs with added Nematoda virus (Orsay NC, n=4) to assess levels of cross-contamination. Both types of NCs were isolated along with the samples and were analyzed in our study.

*Shah* et al.^9^ is a study done in Copenhagen, Denmark that used samples collected as part of the COPSAC2010 cohort^16^. Samples were collected from 647 one-year-old infants one time and isolated along with 8 buffer NCs.

### Sequencing reads: quality control and assembly

We carried out read quality control and assembly, adjusting for study specifications such as the presence of multiple displacement amplification (MDA) during the VLP library construction.

#### Garmaeva et al

Raw reads underwent adapter trimming with the bbduk.sh script from BBTools (v39.01)^17^ using the following flags: ktrim=r k=23 mink=11 hdist=1 tpe tbo. Human read removal and read quality trimming and filtering were performed using kneaddata (v0.10.0)^18^ and human reference genome (GRCh38p13), followed by quality trimming with the option --trimmomatic-options “LEADING:20 TRAILING:20 SLIDINGWINDOW:4:20 MINLEN:50". Quality-trimmed and filtered reads were assembled using SPAdes (v3.15.3) with “--meta” mode^19^. For the one NC sample, SPAdes failed to perform internal sequencing reads processing. We therefore performed read error correction with tadpole.sh (parameters: mode=correct, ecc=t, prefilter=2) from BBTools (v39.01)^17^. Error-corrected and deduplicated reads of NC were assembled with SPAdes (v3.15.3) using the “--–meta” and “--only-assembler” modes^19^.

#### Maqsood et al. and Liang et al

First, we excluded samples SRR8653201, SRR8800143, and SRR8800149 from the Liang et al. study, following the original authors’ recommendations. Raw read quality control and filtering were then performed as described above. Next, to maximize MDA-treated samples’ *de novo* assembly performance, we performed reads deduplication and assembly in the uneven coverage-aware mode^20^. Briefly, read error correction was done with tadpole.sh, as described above, and read deduplication was performed using clumpify.sh (dedupe=t, subs=0) from BBTools (v39.01)^17^ to remove identical sequences. Read assembly was done with SPAdes (v3.15.3) using “--sc” mode^21^.

#### Shah et al

No quality control of the sequencing reads was performed since the deposited samples were already filtered for low-quality and human reads and deduplicated. Assembly was performed with SPAdes (v3.15.3), using “--sc” mode^21^.

### Putative virus sequence prediction from the metaviromes

Putative virus sequences were predicted per sample using assembled contigs >1,000 bp. To maximize the likelihood of virus sequence recognition, we applied four different tools: VirSorter2^22^ (v2.2.4) with the flag “--include-groups “dsDNAphage,RNA,NCLDV,ssDNA,lavidaviridae"”, DeepVirFinder^23^ (v1.0), geNomad^24^ (v1.7.4) in “--end-to-end” mode with enabled score calibration, and VIBRANT^25^ (v1.2.1) with the open reading frames predicted using prodigal-gv (v2.9.0-gv)^24,26^ as an input. All contigs identified as viral by VirSorter2, geNomad, or VIBRANT and DeepVirFinder-identified sequences with a score ≥0.94^27^ were selected for further analysis.

We also attempted to extend identified putative virus sequences using COBRA^28^ (v1.2.3) to recover more complete viruses. To trim host-associated regions from provirus sequences, we subsequently employed geNomad^24^ (v1.7.4) and CheckV (v1.0.1)^29^. Sequences containing direct terminal repeats (DTR) or inverted terminal repeats (ITR) identified by geNomad were exempted from running through CheckV. Genome completeness was estimated for all sequences after prophage-pruning using CheckV.

### vOTU processing and vOTU table creation

971,583 viral contigs were further dereplicated using the MIUVIG recommended cut-offs for the species-level rank: 95% average nucleotide identity (ANI) over 85% alignment fraction (relative to the shorter sequence)^30^, resulting in 307,938 vOTU representatives. To estimate vOTU abundances, we used Bowtie2^31^ (v2.4.5) in ‘end-to-end’ mode to align reads to the vOTU-representative genomes, followed by count table generation using SAMTools^32^ (v1.18) and BEDTools^33^ (v2.30.0). Read counts with a breadth of coverage less than 1×75% of a contig length were set to 0 to remove spurious Bowtie2 alignments^34^. Final read counts were transformed to reads per kilobase per million reads mapped (RPKM) values, which were subsequently used for downstream analyses. We further removed vOTU representatives from the RPKM table if they were shorter than 1,000 bp or had fewer viral genes than host genes based on CheckV assignment. Sequences identified as plasmids by geNomad^24^ (v1.7.4) were also removed from the table. The final RPKM table included 193,970 vOTU representatives detected in 1,368 samples, including 55 NCs.

### vOTU taxonomic profiling and host prediction

We used geNomad^24^ (v1.7.4) taxonomic assignment for the resulting vOTU representatives. Using the taxonomic assignment, we derived information about viral nucleic acid type and predicted host domain (eukaryote or prokaryote) from ICTV^35^. The iPhoP (v1.3.3) framework with database “Aug_2023_pub_rw” was used for host prediction for vOTU representatives^36^. For this analysis, we excluded vOTU representatives identified as eukaryotic viruses via taxonomy assignment. The genus-level host prediction with the top confidence score per vOTU representative of at least 50% genome completeness, as predicted by CheckV, was selected for further analysis. We carried out species-level host prediction for two phages predicted to infect *Burkholderia* using a combination of the iPhoP host genome assignment and blast against nt database (update from 2024/08/14), as BACPHLIP^37^ (v0.9.6) predicted one of the phages to be temperate. Based on the identified ANI of >96% and 76% query coverage, the host was narrowed down to *Burkholderia cepacia* complex species.

### Virus strain reconstruction and comparison

For the 5,635 vOTU representatives present in the biological samples and at least one NC, we reconstructed consensus sequences using inStrain^38^ (v1.9.0). Briefly, the sorted sequence alignment map files of samples and NCs where the vOTU representative of interest was identified were processed with the ‘profile’ module of inStrain in the “--database_mode” with a minimum coverage requirement for variant calling of 1 (“--min_cov 1”). Next, we used the ‘compare’ module of inStrain to estimate the pairwise genome similarity among all reconstructed consensus sequences. Only regions with a minimum of 1X coverage were included in the analysis. Consensus sequence pairs with less than 75% of the genome available for comparison were excluded from the analysis. Population-level ANI (popANI) values were used to compare the similarity between strains belonging to the same vOTU. Virus strains were considered to be shared between samples if their compared genomic regions exhibited ≥ 99.999% popANI^38^.

### Virus sequence decontamination from negative control sequences

To remove virus sequences shared between biological samples and NCs at strain-level, we zeroed out the RPKM values of the relevant vOTU representatives. To account for potential limitations due to varying sequencing depth between samples and NCs, we considered all cases of strain-sharing, including those with less than 75% of the genome available for comparison. For the comparison of strain- and species-level decontamination, vOTUs detected in the NCs and samples from the same study were zeroed out.

### Data visualization

Results were visualized in graphical form using a set of custom R scripts (R v4.2.3), including calls to functions from the package ggplot2^39^ (v3.5.0). All boxplots were prepared using ggplot2 and represent standard Tukey type with interquartile range (IQR, box), median (bar) and Q1 – 1.5 × IQR/Q3 + 1.5 × IQR (whiskers). Phylogenetic trees were built based on the estimated pairwise genome dissimilarity (1 – popANI), using only consensus sequence pairs for which at least 75% of the genome was available for comparison. Hierarchical clustering was applied to the matrices of genome dissimilarity using the function hclust() from the R package stats v.4.2.1^40^. Clustering trees were then converted into phylogenetic trees with the function as.phylo() from R package ape v.5.7.−1^41^. Phylogenetic trees were visualized using the ggtree() function from the R package ggtree^42^.

### Statistics and reproducibility

The current study uses biological samples and NC samples from publicly available datasets. No prior selection was applied to the samples, and no statistical method was used to predetermine sample size. Of the 1,321 biological samples and 55 NCs retrieved, we used 1,313 and 55, respectively, for the statistical analyses.

Statistical analyses were performed using R (v4.2.3). To estimate differences between samples (N=1,313) and NCs (N=55) in features such as number of clean reads, reconstructed contigs, discovered viral sequences, viral richness, and viral diversity, we used linear mixed models (lme4, v1.1-23^43^, and lmerTest, v3.1-3^44^) (Supplementary Data 2–4, 6–7). We then used the same approach to estimate differences between the samples (N=1,254) and NCs (N=37) that had at least one virus detected in viral sequences with at least 50% completeness and viral composition (Supplementary Data 5, 9, and 11). In each comparison, the studies were analyzed both together and separately. When analyzed together, the study and subject group were included as nested random factors. For the biological samples, the subject group was defined as the subject ID to account for the repeated measurements, while NCs were grouped by the source within each study (Supplementary Data 1). When analyzed separately, the subject group was used as a random factor. Garmaeva et al. was excluded from the per-study analysis due to the availability of only one NC. P-values were adjusted using the Benjamini-Hochberg correction method. Viral diversity (Shannon index) was calculated using the “diversity” function in the vegan package v2.6-4^45^.

For the viral composition analysis, the proportions of vOTUs with assigned nucleic acid type and predicted hosts were calculated using only vOTU representatives with at least 50% completeness, based on a binary RPKM table (presence/absence data). These data were log-transformed prior to calculations (Supplementary Data 9, 11).

To assess the similarity between NCs and samples, we calculated Bray-Curtis dissimilarity indices using the “vegdist” function from the vegan package and subtracted from 1. A two-sided Wilcoxon rank sum test, conducted through a permutation approach with n=10,000 iterations, was performed to determine the significance of differences observed between groups (Supplementary Data 12).

The percentage of vOTUs shared between biological samples and NCs was calculated as the proportion of shared vOTUs to the total number of viruses detected per sample. A two-sided Wilcoxon rank sum test, conducted through a permutation approach with n=10,000 iterations, was performed to determine the significance of observed differences in sample vOTU-sharing with NCs from the same study versus NCs from different studies (Supplementary Data 13). To assess the correlation between sample vOTU-sharing with same-study NCs versus with different studies’ NCs, we calculated a study-wise Spearman correlation coefficient (Supplementary Data 14). P-values were adjusted using the Benjamini-Hochberg correction method.

For the analysis relating sample vOTU-sharing with the same study NC to participant type (infant versus mother), we used only biological samples from Garmaeva et al. (N=205) and Maqsood et al. (N=78), as maternal samples were only available in these studies (Supplementary Data 15). For Garmaeva et al., significance was estimated using linear mixed-effects models, with the subject group added as a random factor. For Maqsood et al., we used a linear model because only one timepoint was available per participant. P-values were corrected using the Benjamini-Hochberg method.

For the analysis of vOTU-sharing with same-study NCs over time, we used only the two studies with multiple timepoints per subject (Supplementary Data 16): Garmaeva et al. (N=205) and Liang et al. In Liang et al., only one cohort had multiple timepoints per infant available, so only this cohort’s samples (N=146) were used for the analysis. The RNA samples from Liang et al. were excluded from this analysis. Significance was assessed separately for each study using linear mixed-effects models, with the subject group included as a random factor. P-values were corrected using the Benjamini-Hochberg method.

## Data availability

All data used in this study are publicly available, with sequencing data for Maqsood et al., Liang et al., and Shah et al. accessible via the European Nucleotide Archive (project numbers PRJEB33578, PRJNA524703, and PRJEB46943, respectively). For Garmaeva et al., sequencing data are available in the European Genome-Phenome Archive (EGA) repository (study ID: EGAS00001005969). The source data, including redundant virus sequences and vOTU representatives, along with their metadata, and the database of virus sequences identified in the negative controls (v1.0.0) are available in the FigShare repository under https://doi.org/10.6084/m9.figshare.27170739. Other datasets or databases used in the present study were: Human reference genome GRCh38.p13 [https://www.ncbi.nlm.nih.gov/datasets/genome/GCF_000001405.39/] and iPHoP database (Aug_2023_pub_rw) [https://portal.nersc.gov/cfs/m342/iphop/db/iPHoP.latest_rw.tar.gz].

## Code availability

The code used in this study can be found at: https://github.com/GRONINGEN-MICROBIOME-CENTRE/NCP_VLP_project

## Acknowledgements

We thank Kate Mc Intyre for editing the manuscript. We also thank the Genomics Coordination Center and the Center for Information Technology of the University of Groningen for their support and for providing access to the Gearshift and Hábrók high-performance computing clusters. SG was supported by a scholarship from the Graduate School of Medical Sciences, University of Groningen. SG was awarded a de Cock-Hadders Stichting grant (2021-08). Furthermore, this project was funded by the Netherlands Organisation for Scientific Research (NWO): NWO Gravitation Exposome-NL (024.004.017) to AK and AZ and an NWO-VICI VI.C.232.074 to AZ. AZ was also supported by the EU Horizon Europe Program grant INITIALISE (101094099). AK was supported by ZonMW ME/CFS grant 10091012110017.

## Contributions

S.G. designed the study. N.K. gathered and prepared the data. N.K., A.K., and S.G. analyzed the data. N.K., A.Z., and S.G. wrote the manuscript. A.K. provided advice for statistical methods. N.K., A.K., A.Z., and S.G. critically reviewed the manuscript.

