## Supplementary figures for "Negativeome in Early-Life Virome Studies: Characterization and Decontamination"

N. Kuzub et al.

Supplementary Figure 1: Study design and sample distribution per dataset.

Supplementary Figure 2: Characteristics of viral sequences used for species-level dereplication.

Supplementary Figure 3: Comparison of genomic and ecological features between biological samples and NCs.

Supplementary Figure 4: Percentage of vOTUs shared with negative controls (NCs) across samples, studies, and age groups.

**Supplementary Figures**

| Sample  type | Study | NC source / sample timepoint | N  extracted | N deposited | N  after read QC | N  with non-zero richness | N  sharing vOTUs to NCs |
| --- | --- | --- | --- | --- | --- | --- | --- |
| 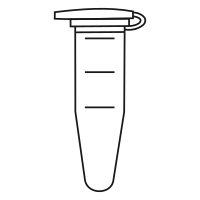NCs | Garmaeva et al. | Buffer | 4 | 1 | 1 | 1 | 1 |
|  | Liang et al. | Buffer; Tube; Diaper;  MDNC | 38 | 38 | 38 | 20 | 20 |
|  | Maqsood et al. | Buffer; Orsay | 8 | 8 | 8 | 8 | 8 |
|  | Shah et al. | Buffer | 8 | 8 | 8 | 8 | 8 |
| 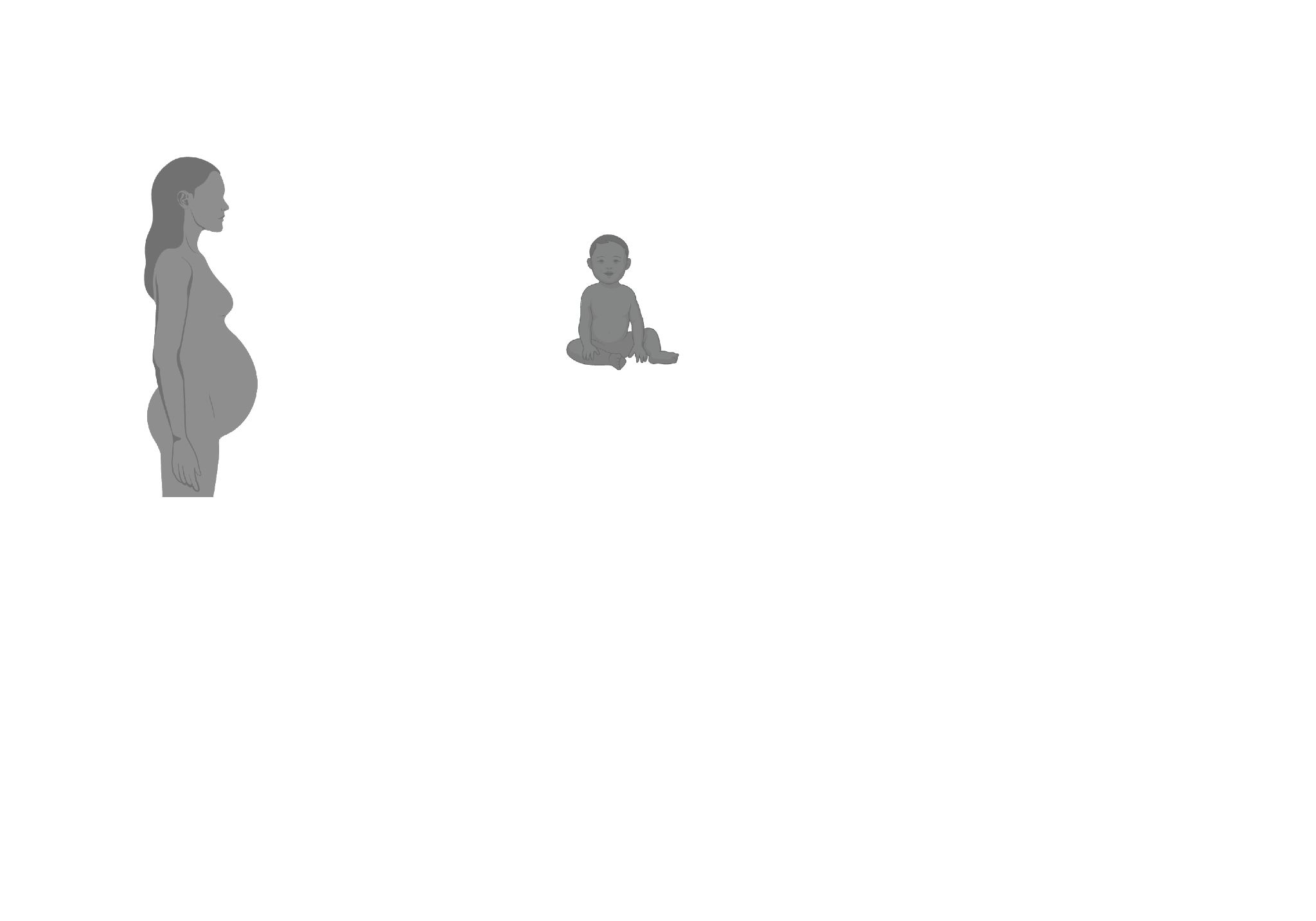Infants | Garmaeva et al. | M1; M2; M3; M6; 12 | 129 | 86 | 86 | 86 | 72 |
|  | Liang et al. | M0; M1; M4; Y2-5 | 394 | 391 | 383 | 324 | 137 |
|  | Maqsood et al. | M0 | 56 | 51 | 51 | 51 | 42 |
|  | Shah et al. | M12 | 660 | 647 | 647 | 647 | 620 |
| 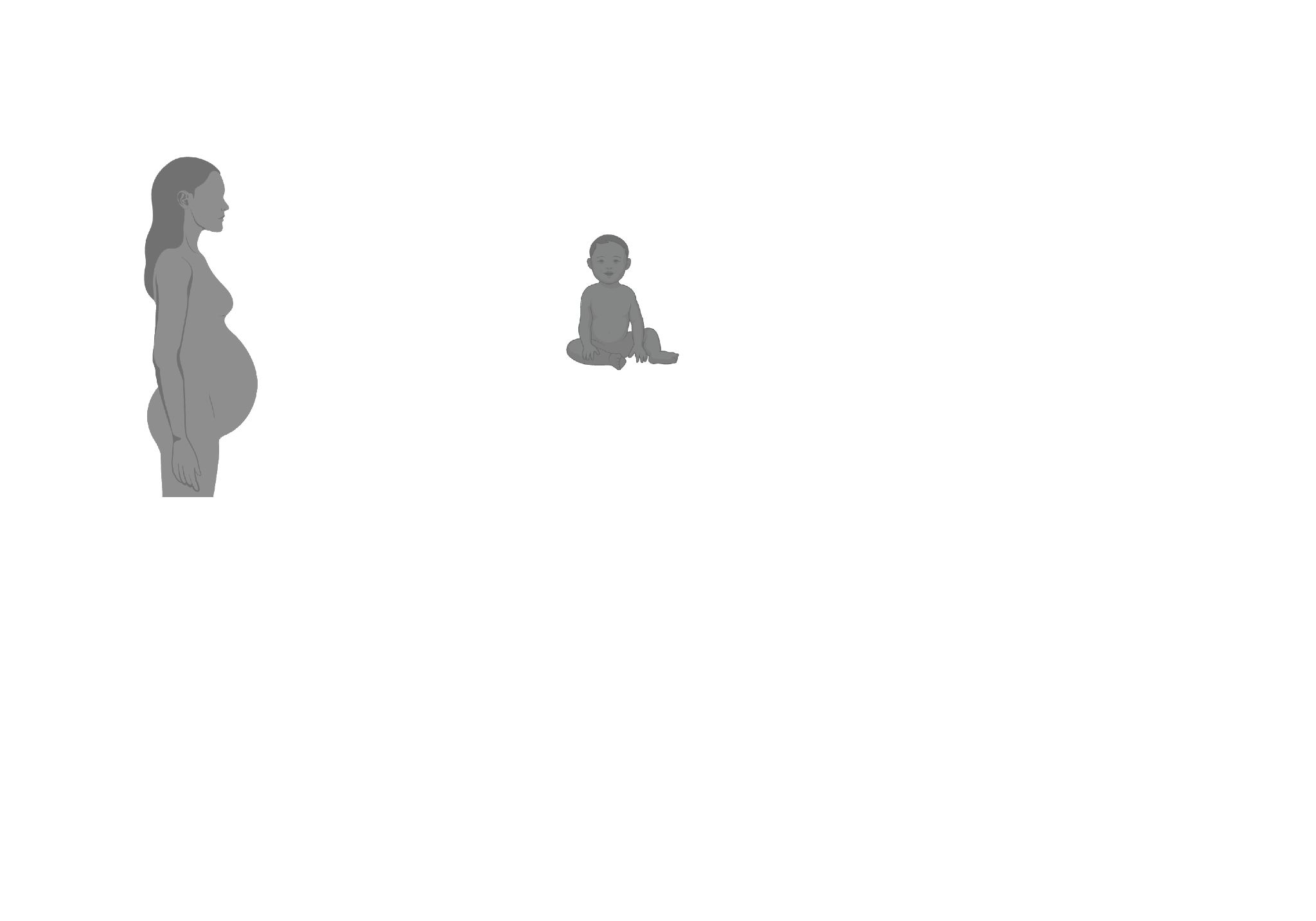Mothers | Garmaeva et al. | Mtrim3; M0; M1; M2; M3 | 223 | 119 | 119 | 119 | 71 |
|  | Maqsood et al. | M0 | 28 | 27 | 27 | 27 | 2 |

**Supplementary Figure 1. Study design and sample distribution per dataset.** The third column provides the source of negative controls (NCs) for each study, while for biological samples, it lists the timepoints included per study. Timepoints are represented by the abbreviations M0, M1, M2, M3, M4, M6, and M12, corresponding to the infant's age in months at the time of sampling (M = month). Additionally, "Mtrim3" denotes samples collected during the mother's third trimester of pregnancy, and "Y2-5" indicates samples collected from infants aged 2 to 5 years. Detailed information on the number of samples per timepoint for both infants and mothers can be found in Supplementary Data 1. Columns 4-8 provide further details: the number of samples that were extracted using viral-like particle enrichment protocol (N extracted); the number of samples deposited to the archives (N deposited); the number of samples with non-zero clean reads following read quality control (N after read QC); the number of samples where at least one vOTUr was identified (N with non-zero richness); the number of samples that shared at least one vOTUr with the NCs (N sharing vOTUs to NCs).

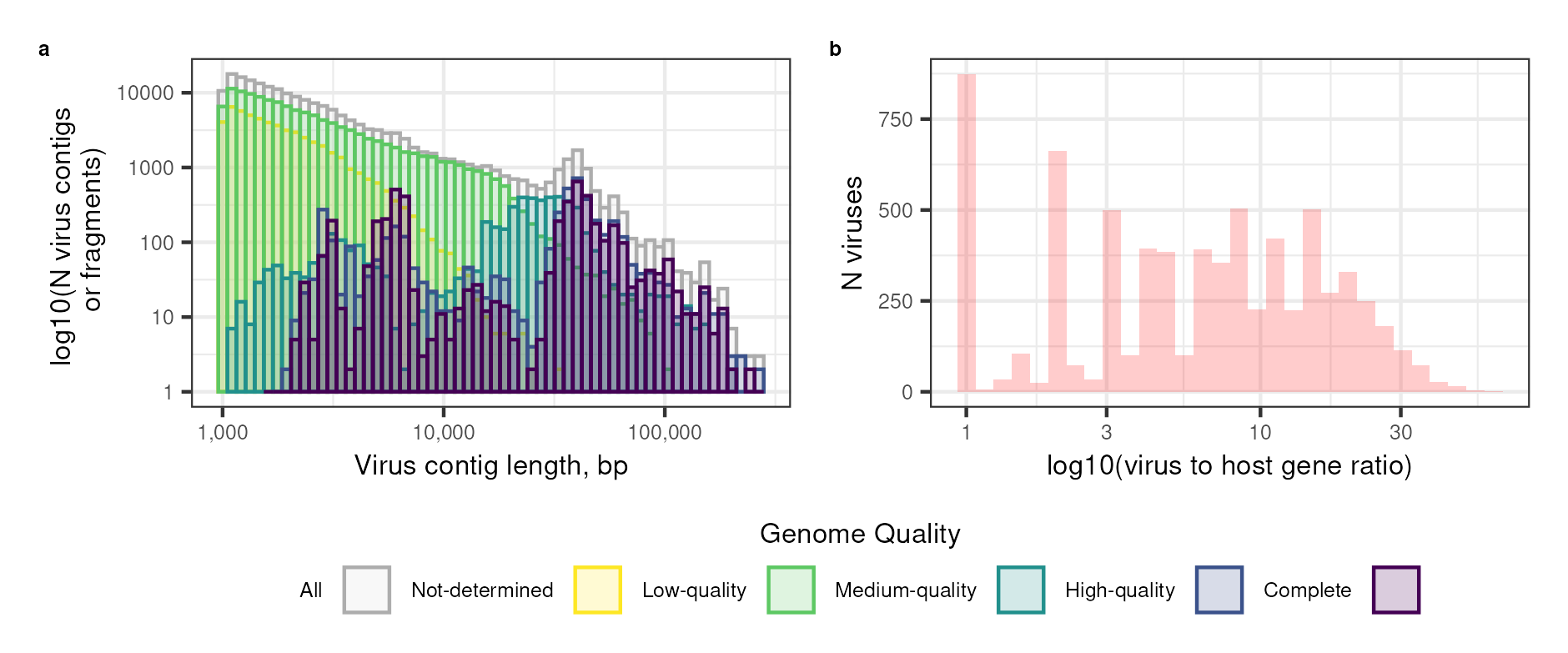

**Supplementary Figure 2. Characteristics of viral sequences used for species-level dereplication. a.** The length distribution of viral genomes and genome fractions coloured by genome quality estimated by CheckV. The Y-axis is displayed on a logarithmic scale. **b.** Distribution of viral genomes and genome fractions based on the virus-to-host gene ratio. In **a-b,** the data presented reflects the virus genomes and genome fractions that fulfilled the following criteria: length > 1kbp, a higher number of viral genes compared to host genes, and no plasmid sequences.

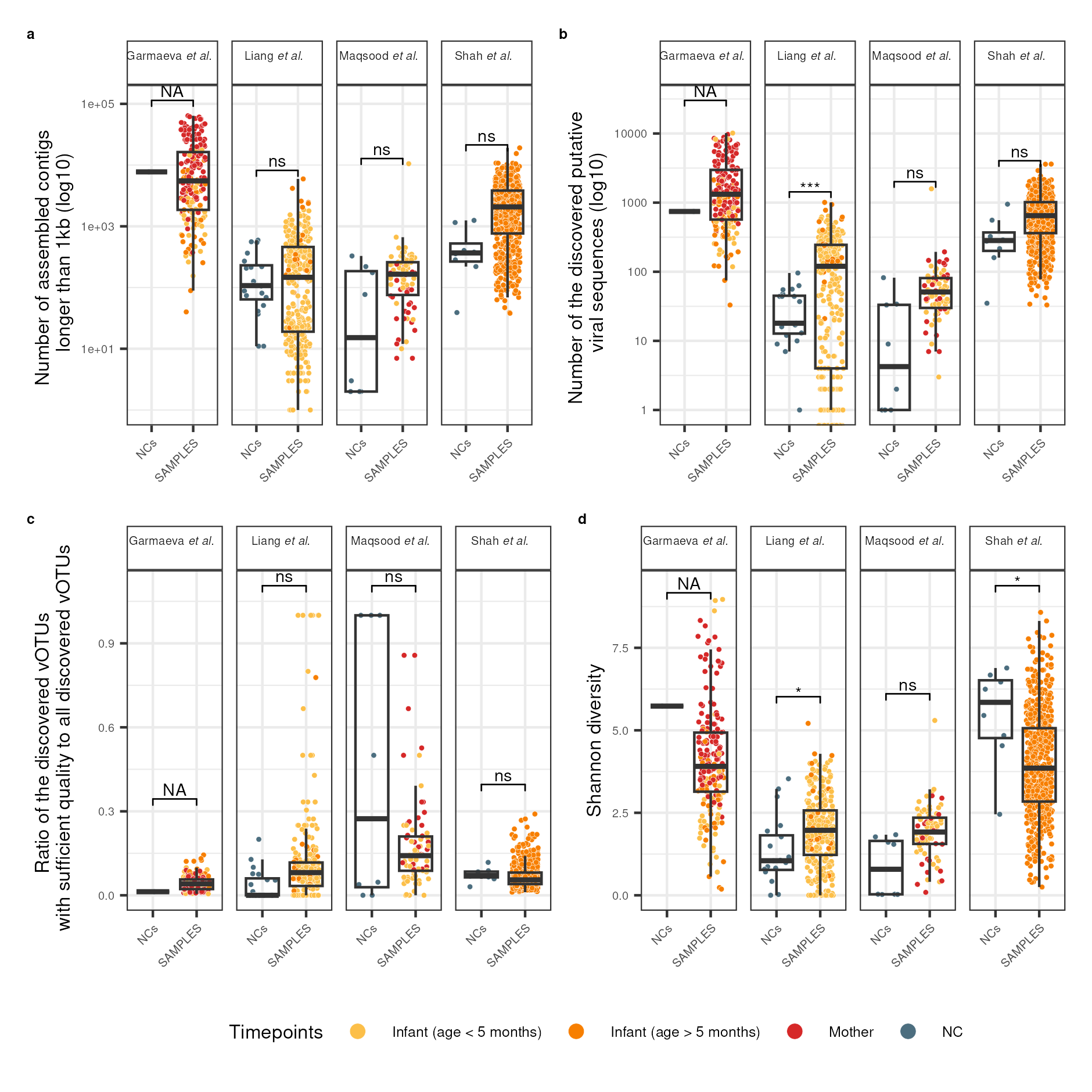

**Supplementary Figure 3. Comparison of genomic and ecological features between biological samples and NCs. a.** Number of assembled contigs longer than 1kb in NCs compared to biological samples. **b.** Number of the discovered putative viral sequences in NCs vs samples. **c.** Ratio of the discovered putative viral sequences of sufficient quality to all putative viral sequences in NCs vs samples. Sufficient quality indicates viral sequences of at least 50% completeness as assessed by CheckV. **d.** Shannon diversity in NCs vs samples. In **a-d,** each sample is a dot, and the dot colour represents the age: infant samples (age < 5 months) in yellow, infant samples (age > 5 months) in orange, maternal samples in red, and NCs in dark blue. Boxplots visualise the median, hinges (25th and 75th percentiles), and whiskers extending up to 1.5 times the interquartile range from the hinges. **a-c** are shown in the logarithmic scale.

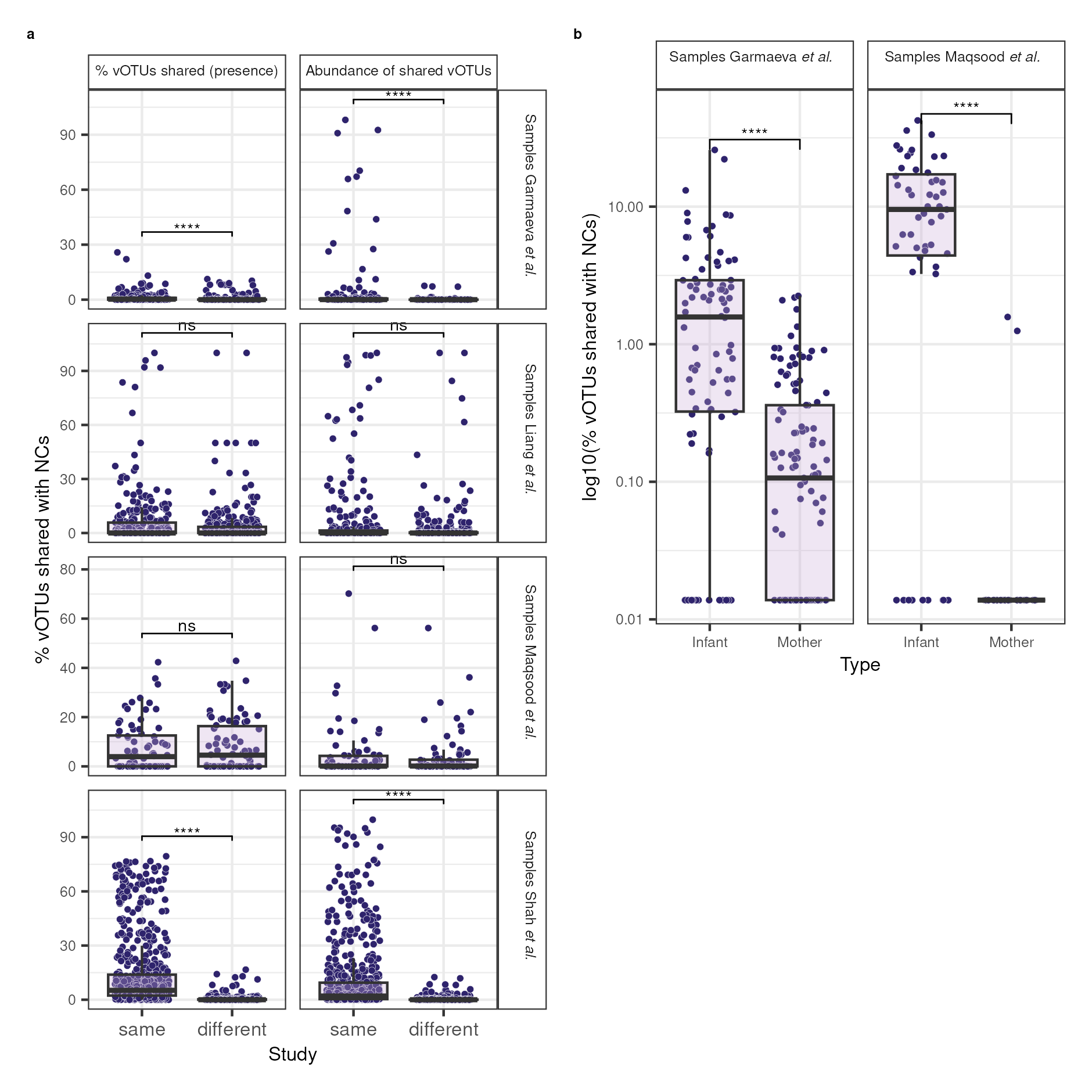

**Supplementary Figure 4. Percentage of vOTUs shared with negative controls (NCs) across samples, studies, and age groups. a.** Percentage and abundance of vOTUs shared with NCs from the same study compared to the percentage and abundance of vOTUs shared with NCs from different studies combined. The percentage of vOTUs shared with NCs is calculated as the number of vOTUs shared with NCs divided by the total sample richness. **b.** Percentage of vOTUs shared with NCs from the same study in maternal samples compared to the infant samples. Y-axis is shown in the logarithmic scale. In **a-b**, boxplots show the median, hinges (25th and 75th percentiles), and whiskers extending up to 1.5 times the interquartile range from the hinges.
